# *Plasmodium falciparum* growth is regulated by Sphingosine 1 phosphate produced by Host Erythrocyte Membrane Sphingosine kinase 1

**DOI:** 10.1101/756502

**Authors:** Raj Kumar Sah, Monika Saini, Soumya Pati, Shailja Singh

**Author notes:** Correspondence: Shailja Singh.

## Abstract

Sphingosine-1-phosphate (S1P) a bioactive lipid is produced in its primary reservoir, erythrocytes by an enzyme Sphingosine kinase-1 (SphK-1). The activation of such kinases and the subsequent S1P generation and secretion in the blood serum represent a major regulator of many cellular signaling cascades. Orthologue of sphingosine kinases 1 and 2 (SphK-1 and 2) that catalyze the phosphorylation of sphingosine generating S1P are not present in malaria parasite. The malaria parasite, *Plasmodium falciparum*, is an intracellular obligatory organism that reside in the human erythrocyte during its blood stage life cycle and orchestrates many metabolic interactions with host for its survival. Given the regulatory role of S1P, we targeted host SphK-1 by a generic pharmacological inhibitor N,N-Dimethyl-sphingosine (DMS) and analyzed growth of intra-erythrocytic parasite. We found that reducing S1P levels by inhibiting host SphK-1 activity led to halted parasite growth and ultimately cell death. Reduced intracellular S1P levels were attributed to decreased glycolysis marked by the low uptake of glucose by parasite and by less production of lactate, a byproduct of glycolysis. Reduced glycolysis was mediated by decrease translocation of the glycolytic enzyme, Glyceraldehyde 3-phosphate dehydrogenase (GAPDH) to the cytosol of infected erythrocytes and cell death. Knocking down of erythrocyte SphK-1 is not lethal to the host and being a host encoded enzyme, targeting it with safe and specific drugs will not lead to the problem of resistance; thus, SphK-1 represents a potent target for the development of therapeutics against intra-erythrocytic *P. falciparum*.

**Author Summary:** Erythrocytes membrane enzyme Sphingosine kinase-1 (SphK-1) produces Sphingosine-1-phosphate (S1P) a bioactive lipid by phosphorylation of Sphingosine (Sph). S1P generated by activation of SphK is prosurvival signal and regulate cell growth. The malaria parasite, *Plasmodium falciparum*, is an intracellular obligatory pathogen that reside in erythrocyte during its blood stage life cycle and orchestrates many metabolic interactions with its host erythrocytes for survival. Orthologue of SphK-1/ 2 are not present in malaria parasite, therefore treatment with SphK inhibitor targeted host SphK-1 and led to reduced S1P level. The reduction in host S1P led to halted parasite growth and cell death. Furthermore, reduced erythrocyte S1P levels led to decreased glycolysis marked by the low uptake of glucose by parasite and by less production of lactate. Erythrocyte SphK-1 being a host encoded enzyme, is resistance safe and represents a potent target for the development of therapeutics against intra-erythrocytic *P. falciparum*.

## Introduction

An estimate of 219 million malaria cases and 435,000 related deaths were reported in 2017 worldwide according to World Health Organization[1]. The decrease in efficiency of current anti-malarial agents including artemisinin, quinine, chloroquine, piperaquine, mefloquine and their derivatives in malaria affected regions of the world has significantly increased the cost and complexity of curing malaria[2],[3],[4]. The limited number of available anti-malarial drugs and parasitic resistance to almost every available chemical therapy continues to spur the search for novel approaches. The next generation of anti-malarials are in pipeline and holds a great promise as they target novel parasite encoded enzymes and molecular pathways[5],[6]. However, since these drugs target molecules that are under genetic control of the parasite, resistance against them can be developed in the long run[7],[8]. To deal with the nuisance of drug resistance, targeting host encoded proteins that are indispensable for parasite growth and survival would be an ideal situation. Since, the chances of development of resistance against non-parasite targets are very bleak, host targeted drug development will be a classical addition to the field of drug discovery in malaria.

*P. falciparum* being an intracellular obligatory parasite exploits the host’s resources and pathways for its survival and growth. For example, parasite feeds on erythrocytic hemoglobin content during the asexual blood stage development. The breakdown of hemoglobin provides amino acids for its growth and maturation[9]. Another important example is the metabolic pathway of glycolysis. It is well known that the parasite infected erythrocytes utilize glucose at a much higher rate than the normal parasite uninfected erythrocytes[10],[11]. During the intra-erythrocytic growth phase, *Plasmodium* lacks a functional tricarboxylic acid (TCA) cycle (also known as Krebs cycle) and is therefore, dependent on glycolysis for its energy requirements[12]. For glycolysis, the parasite makes use of pre-existing pools of host glucose as well as imports it by expressing glucose transporters on the surface of the host erythrocytes[13]. The parasite’s sole dependence on glycolysis for energy needs makes it a potential target for anti-malarial chemotherapies. However, the regulation of glycolysis by parasites in infected erythrocytes is not very clear[14],[15],[16].

A recent study demonstrates that intracellular S1P facilitates glycolysis in erythrocytes in response to hypoxia[17]. This biolipid is involved in various other biological processes including immune response[18], bone marrow cells trafficking [19], vascular integrity[20], cell survival and proliferation[21]. Given the diverse roles of S1P, various cell types have been identified as production and store house of S1P including erythrocytes, endothelial cells, thrombocytes, mast cells, and macrophages[22],[23]. However, erythrocytes have been considered as the main repository for S1P in the blood plasma[22],[24]. Several reasons have been held responsible for the elevated S1P content in these cells including high sphingosine kinase (SphK) activity, lack of S1P degrading enzymes (S1P lyase and S1P phosphohydrolase) and its capability to import sphingosine from extracellular environment[25],[26]. Surprisingly, the role of multi-faceted S1P in erythrocytes and what affect it exerts on the physiology of the cells remains an unexplored mystery. S1P is produced through hydrolysis of sphingomyelin to ceramide by sphingomyelinases, followed by sphingosine synthesis from ceramide via the action of cermidases, and finally phosphorylation of sphingosine via sphingosine kinases (SphK) produces S1P[27]. S1P is basically a phosphorylated product of sphingosine produced by kinases, sphingosine kinase-1 and -2 (SphK-1 and SphK-2)[28]. While SphK-1 is localized to the cytosol, SphK-2 resides in the nucleus[29],[30]. Erythrocytes harbor only SphK-1 which is the main enzyme responsible for the production of S1P in them[22],[24]. Phosphorylation of host SphK-1 acts as a key regulating factor for its activity. SphK-1 selectively binds to phosphatidylserine in the membrane and phosphorylation at serine 225 is essential for its increased selective membrane binding capability[28]. Intracellular erythrocytic S1P binds to deoxygenated-hemoglobin, translocates to the membrane and mediates the release of the glycolytic enzyme, GAPDH, which in turn regulates the erythrocytic glycolysis pathway[17]. Apart from being synthesized within the erythrocytes, S1P gets secreted out in the blood plasma via a specific S1P transporter major facilitator superfamily transporter (Mfsd2b)[31].

With this background, we aimed to study the role of host SphK-1 in regulation of parasite growth inside the host erythrocytes. We demonstrated inhibition of host SphK-1 by N,N-Dimethylsphingosine (DMS) lowered S1P levels in the host. The Inhibition of host SphK-1 led to the reduction in glycolysis and cell death of parasite. The Inhibition of SphK-1 was found to be associated with reduction in cytosolic presence of glycolytic enzyme GAPDH. Glycolysis in normal erythrocytes is mainly regulated by an essential cytosolic enzyme, GAPDH, which remains bound to the membrane unless required during glycolysis[32]. Thus, reduction of cytosolic GAPDH leads to lowered activity in parasite-infected erythrocytes can be an explanation for parasite death. Since deletion of host SphK-1 does not have any adverse effect on host, we advocate the targeting of host SphK to kill parasite for the development of potent anti-malarial drugs. Also, since it is a host-encoded enzyme, the possibility of the resistance development by the parasites is diminished.

## Materials and methods

### Cultivation of *P. falciparum*–infected erythrocytes

*P. falciparum* 3D7 strain was cultured in RPMI 1640 (Invitrogen, Carlsbad, CA, USA) supplemented with 27.2 mg/L hypoxanthine (Sigma-Aldrich, St. Louis, MO, USA), 2 gm/L sodium bicarbonate (Sigma-Aldrich, St. Louis, MO, USA) and 0.5 gm/L AlbuMax I (Gibco, Grand Island, NY, USA) using O+ human erythrocytes, under mixed gas environment (5% O_2_, 5% CO_2_ and 90% N_2_) as described previously[33]. For the assessment of half-maximal drug concentration for inhibition of malaria survival (IC_50_) values, synchronized *P. falciparum* infected erythrocyte cultures were used at late-ring or early trophozoite stage (18-24 hours post infection (hpi)) at a parasitemia of 1%. Where indicated, cultures were treated with the DMS inhibitor (Sigma-Aldrich, St. Louis, MO, USA) at concentrations ranging from 0 to 40 µM. Untreated controls were cultured in parallel under the same conditions and processed identically. To assess total parasitemia, cultures were collected at the indicated times and freeze-thawed. The lysates were processed for SYBR-green staining (Thermo Fisher Scientific, Waltham, Massachusetts, US). Briefly, equal volumes of lysis buffer containing 20 mM tris (pH 7.5), 5 mM EDTA, 0.008% saponin (*w/v*), and 0.08% triton X-100 (*v*/*v*) was added to the lysed parasites and incubated for 3 hours at 37°C with 1X SYBR-green dye. Fluorescence after the SYBR-green assay was recorded in a multimode plate reader (Bio□Rad) at an excitation and emission wavelength of 485 nm and 530 nm, respectively. Percent growth inhibition was calculated using the following formula: % Growth Inhibition = {(Control - Treated)/ (Control)* 100}. The effect of DMS was tested on progression of the parasites treated at ring stage and monitored till different developmental asexual stages, namely, rings, trophozoites, schizonts and release of merozoites from schizonts. Tightly synchronized ring stage parasite cultures treated with 10 µM DMS inhibitor or solvent as control were incubated for 0, 18, 34, 45 hours blood stage asexual cycle to monitor the progression at each stage. Morphological analysis and counting (∼3,000 cells/Giemsa-stained slides in duplicate) were done at each of these stages to monitor the parasite’s progression.

### Analysis of morphological changes in infected and uninfected erythrocytes by Scanning Electron Microscopy (SEM) after DMS treatment

The infected and uninfected erythrocytes were treated with DMS (10 µM) for a period of 5 hours. Following incubation erythrocyte and infected erythrocyte were washed three times in sterile PBS. The samples were then fixed by 2.5% glutaraldehyde in 1X PBS (pH 7.4) with 2% formaldehyde for a period of 30 minutes. Post fixation, samples were rinsed thrice with 1X PBS and dehydrated in absolute ethanol series (ethanolic dehydration), using a standard protocol. Samples were then completely dried, coated with gold, and observed under the scanning electron microscope.

### Extraction of Lipids and Sphingosine-1-Phosphate Measurement

Equal number of synchronous parasitized erythrocytes at 7-8% parasitaemia, and uninfected erythrocytes both in presence and absence of DMS for a period of 5 hours were used for further experiments. Collected cell pellets and supernatant were employed for lipid extraction as reported previously [34]. Pellets were resuspended in 100 µl H_2_O and transferred to 900 µl methanol. Whereas, for quantification of S1P in supernatant, 1:15 ratio of methanol was used. After vortexing and centrifugation at 10,000 ×*g* for 5 minutes at RT, methanol extracts were removed to a new glass tube. After evaporation by N_2_, dried lipids were resuspended in 200 µl methanol. Extracted lipid samples were subjected to liquid-chromatography mass-spectrometry (LC/MS) analysis. We used a Waters Acquity H-Class UPLC-system (Waters, Milford, MA, USA). Chromatographic separation was achieved on an Acquity BEH C18, 1.7 µm, 75 × 2.1 mm column (Waters, Manchester, UK). Mass spectrometry was performed in negative electrospray mode using a high-resolution mass spectrometer synapt G2 S HDMS (Waters, Manchester, UK) with a TOF-detector with linear dynamic range of at least 5000:1. The mass spectra were acquired over the range of 100–1000 Da with a spectral acquisition rate of 0.1 seconds per spectrum.

For S1P measurement by enzyme-linked immunosorbent assay (ELISA) (MyBioSource, San Diego, USA) method, parasite infected and uninfected erythrocytes were treated with 10 µM DMS for 5 hours at 37°C. Collected supernatant was used to measure extracellular S1P, whereas, the lysed erythrocytes were used to measure intracellular S1P level. The samples were added to the micro ELISA plate wells separately, pre-coated with S1P-specific antibody for 90 minutes at 37°C followed by probing with a biotinylated detection antibody specific to human S1P and incubation for 1 hour at 37°C. After washing, Avidin-horseradish peroxidase (HRP) conjugate was added successively to each microplate well and incubated for 30 minutes at 37°C. The wells were washed to remove the unbound components followed by addition of substrate solution to each well. Only those wells that contained S1P, biotinylated detection antibody and Avidin-HRP conjugate appeared blue in color. The enzyme-substrate reaction was terminated by addition of stop solution turning the reaction color to yellow. The optical density (O.D.) proportional to the S1P level was measured spectrophotometrically at a wavelength of 450 nm.

### Fluorescence microscopy for uptake of NBD-Sphingosine in infected erythrocyte

Washed infected erythrocyte (1 × 10^8^/mL) were incubated with 1 µM N,N Dimethyl sphingosine (Sigma Aldrich) or 1 µM omega (7-nitro-2-1, 3-benzoxadiazol-4-yl)(2S,3R,4E)-2-amino octadec-4-ene-1,3-diol (NBD–sphingosine; Avanti Polar Lipids) for 15 min at 37 °C. The infected erythrocytes were pelleted at 550 ×g for 5 min and resuspended in fresh incomplete RPMI media. Approximately 100 µL of the samples was placed onto a glass bottom petri dish. The cells were then allowed to settle at RT for 5 min and were viewed using a confocal Nikon Ti2 microscope equipped with a 100× oil objective (Melville, NY). Digital images were captured. Further, images were processed via NIS-Elements software.

### Immunofluorescence assay and immunoblotting for SphK-1 and GAPDH in erythrocytes

For immunofluorescence assays, thin smears of schizont or mixed stage parasites treated or untreated with DMS were made on glass slides, air dried and fixed with methanol (ice cold) for 30 minutes at −20°C. Smears were blocked with 3% (*w*/*v*) bovine serum albumin (BSA) in phosphate buffer saline (PBS) blocking buffer (pH 7.4) for 30 minutes at room temperature (RT). Slides were probed with anti-SphK-1 (Invitrogen, Carlsbad, CA, USA, 1:1000) rabbit and anti-GAPDH (Invitrogen, Carlsbad, CA, USA, 1:500) mouse antibodies in blocking buffer at RT for 1 hour. After washing, slides were incubated with Alexa Fluor 594 conjugated goat anti-rabbit IgG (Molecular Probes, USA, 1:500) and Alexa Fluor 488 conjugated goat anti-mouse IgG (Molecular Probes, USA, 1:500) at RT for 1 hour. After washing, the slides were mounted in ProLong Gold antifade reagent (Invitrogen, Carlsbad, CA, USA), viewed on a Nikon A1-R confocal microscope and Olympus confocal microscopy. Further, images were processed via NIS-Elements software.

### Immunoblotting for SphK-1 and GAPDH in erythrocytes

Parasite infected and uninfected erythrocytes in the presence or absence of DMS were lysed by freeze-thawing in 10 volumes of 5 mmol/L cold phosphate buffer (pH 8.0) with 1X protease inhibitor cocktail (PIC), vortexed and erythrocyte membrane pellets were separated from cytosolic fraction by centrifugation at 20,000 ×*g* for 20 minutes at 4°C. The fractionated pellets were washed ten times with phosphate buffer to obtain ghost erythrocytes. Total cell pellet of parasite infected and uninfected erythrocytes were re-suspended in RIPA buffer [100 mM phosphate buffer pH 7.2, 150 mM NaCl, 1% NP-40, 0.5% sodium deoxycholate, 0.1% SDS, 50 mM EDTA and 1X PIC. Equal amounts of each sample were boiled with 2X Laemmli buffer, separated on a 10% polyacrylamide gel, transferred onto the PVDF membranes (Millipore) followed by blocking with 5% skim milk blocking buffer for 1 hour at 4°C. After washing, blots were incubated for 1 hour with anti-SphK-1 (1:3000), anti-Phospho-SphK-1 (Ser225) (Invitrogen, Carlsbad, CA, USA, 1:1000) rabbit and anti-GAPDH (Invitrogen, Carlsbad, CA, USA, 1:10,000) mouse antibodies in blocking buffer. Later, the blots were washed and incubated for 1 hour with appropriate secondary antibodies anti-rabbit and anti-mouse (1:10,000) conjugated to HRP. Immunoblotted proteins were visualized by using the Clarity Western ECL substrate (Bio-Rad).

### Glycolysis estimation by lactate level measurement and glucose uptake

Glycolysis estimation was performed using lactate assay kit (Sigma-Aldrich, St. Louis, MO, USA). Parasite infected and uninfected erythrocytes were resuspended in cRPMI containing 10 µM DMS and incubated for 3 hours at 37°C. The supernatant was separated by centrifugation at 5000 ×*g* for 10 minutes at RT and lactate levels were measured in supernatant and lysed erythrocytes according to the manufacturer’s protocol. Glucose uptake into the parasites was quantified with 2-(*N*-(7 Nitrobenz-2-oxa-1, 3-diazol-4-yl) Amino)-2-Deoxyglucose (2-NBDG) (Sigma-Aldrich, St. Louis, MO, USA) via fluorescence labelling and flow cytometry. 2-NBDG is a fluorescent DI glucose derivative and acts as a tracer. Synchronized trophozoite stage parasites were treated with 10 µM DMS and incubated for 3 hours at 37°C. The parasite medium was replaced with RPMI without glucose supplemented with 0.3 mM 2□NBDG followed by incubation for 20 minutes at 37°C to allow uptake of the glucose analogue. Cells were viewed on a Nikon A1-R confocal microscope (Nikon, Tokyo, Japan) and further, images were processed via NIS-Elements software. For flow cytometry, analysis was done on BD LSR Fortessa flow (Franklin Lakes, NJ) and FlowJo software.

### Statistical Analysis

The data for all the assays are expressed as the mean ± standard deviation (SD) of three independent experiments done in triplicates.

## RESULTS

### Inhibition of host SphK-1 activity by specific inhibitor DMS decreases both intracellular and extracellular S1P levels

Physiologically, the main reservoir of circulating S1P is erythrocytes, which are rich in SphK-1 while devoid of S1P lyase and sphingolipid transporter 2 (SPNS2)[25],[26]. Hence, modulation of SphK-1 level could mediate change in the levels of circulating or intracellular S1P concentration. Therefore, we determined the level of SphK-1 in parasite infected and uninfected erythrocytes by immunolabelling and immunoblotting. For immunolabelling, mixed stage parasites were probed with anti-SphK-1 antibody. Significant reduction in SphK-1 levels was found after treatment with DMS in parasite infected erythrocytes as compared to control by two different technique. (Fig. 1a). To further understand, if host Sphk-1 activity might be governed by phosphorylation, we have evaluated the phosphorylation status of host SphK-1 in presence of its specific inhibitor (DMS), by probing lysate of infected erythrocytes with specific anti-SphK-1 and Phospho-SphK-1 (Ser225) antibodies. Whereas, GAPDH was used as a loading control. In line with the previous result, host SphK-1 level was found to be reduced in parasite infected erythrocytes and uninfected erythrocyte treated with DMS as compared to the untreated (Fig. 1b). Moreover, the phosphorylated form of host SphK-1 (pSphK-1) was significantly reduced in parasite infected erythrocytes and uninfected erythrocyte following DMS treatment. The band intensities were quantified to confirm the relative changes in both level of SphK-1 and phosphorylation status (Fig. 1b).

**Figure 1.**
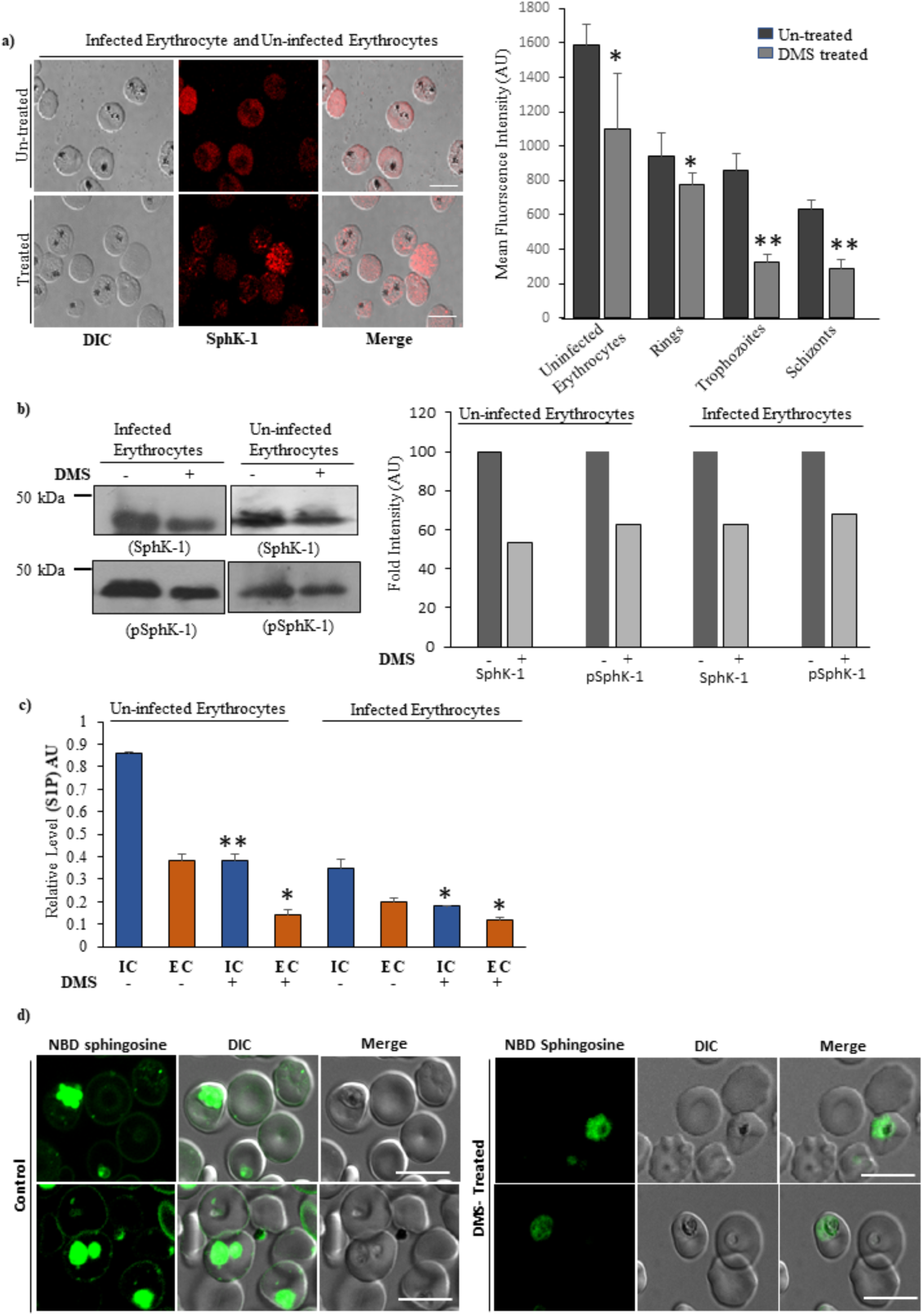
Inhibition of host SphK-1 by specific inhibitor DMS causes decrease in SphK-1 protein level and S1P levels. **a)** Confocal micrographs demonstrate altered host SphK-1 level in mixed-stage parasite culture following treatment with DMS (10 µM). Bar graph denotes the differential mean fluorescence intensity (MFI) denoting host SphK-1 level in infected and uninfected erythrocytes following DMS treatment. **b**) Evaluation of level and phosphorylation status of SphK-1 by immunoblotting. Total cell lysates probed with GAPDH was used as a loading control. The graph represents fold change in the band intensity in individual lanes. **c**) Bar graph depicts ELISA-based S1P quantification in IC and EC microenvironments of infected and uninfected erythrocytes. **d)** NBD-Sphingosine-1-phosphate is localized to membrane and parasite. Infected erythrocyte were resuspended in buffer containing fatty acid-free BSA (0.1% (w/v)) and incubated with DMS (10 µM) or NBD-sphingosine (1 µM) for at least 15 min at 37 °C. The Infected erythrocytes were imaged by fluorescence using a 100× oil objective on a Nikon Ti2 microscope.

Similar results were obtained in a study done parallelly wherein, host SphK-1 level and activity was found to be regulated upon parasite infection to human erythrocytes (manuscript in review with Frontiers). Further, to determine whether host SphK-1 inhibition during parasite infection poses any alteration in S1P levels, we detected the relative levels of S1P in intracellular (**IC**) and extracellular (**EC**) milieu of the erythrocytes through LC/MS analysis and ELISA based kit. Lipid extracts were prepared from *P. falciparum* infected and/or uninfected erythrocytes in the presence or absence of 10 µM DMS and samples were then subjected to LC/MS analysis[34]. A characteristic peak at position ∼ 378.16 in MS spectra was detected for S1P in all the experiments (**Supplementary Figure 1, 2 (A-H)**). Interestingly, the S1P level in DMS-treated infected erythrocytes was almost non-detectable, while the untreated infected erythrocytes demonstrated drastic reduction in S1P levels in both in IC and EC microenvironments. While DMS-treated healthy erythrocytes showed significant reduction in S1P levels from both IC and EC (**Supplementary Figure 1)**. In order to validate the regulation of S1P levels by DMS, ELISA-based quantification was performed. The findings revealed significant down regulation in S1P levels in DMS-treated infected erythrocytes, as compared to untreated infected erythrocytes. Whereas, DMS treatment led to ∼50% reduction in both EC and IC profiles of healthy erythrocytes, when compared to untreated healthy controls (Fig. 1c). These results implicated reduced SphK-1 activity in infected erythrocytes congruent to the effect observed in the uninfected cells on DMS treatment.

Further, we examined whether DMS affect the uptake of NBD-sphingosine or not. For this the infected erythrocyte were incubated with NBD-sphingosine, when NBD-sphingosine added to erythrocyte it incorporated on erythrocyte membrane and get phosphorylated by SPHK-1 into NBD-S1P. phosphorylate by SphK-1 and convert into NBD sphingosine 1 phosphate.[35] The result shows that incubation of infected erythrocyte with NBD-sphingosine it get incorporated into the erythrocyte membrane and parasite and shows fluorescence which suggest the uptake of NBD sphingosine. To further investigate whether the infected RBCs in presence of DMS shows the same phenomenon we incubate the infected RBCs with NBD-sphingosine and DMS. After treatment incubation was done for 15min at 37 and we found that in treated sample NBD-sphingosine get less incorporated in parasite as well as on erythrocyte membrane as compared to the control one. (Fig. 1d).

This data suggests that inhibition of Sphk-1 by DMS causes less uptake of NBP-sphingosine as well as less production of S1P which is supported via LC-MS profiling

### Inhibition of erythrocytic SphK-1 triggers parasite death

To decipher the link between host SphK1 activity and parasite growth, we have elucidated the effect of DMS on synchronized parasite cultures at ring stage. DMS treatment was done in an increasing manner (0, 5, 10, 20, 30 and 40 µM) for 48 hours. After completion of one asexual life cycle of the parasite, SYBR-green was added for assessing the percentage inhibition by DMS. The data revealed a moderate to potent anti-malarial activity of DMS with a significant reduction in the parasite load, with IC_50_ value attended at 14 µM (Fig. 2a). To examine its impact on stage specific inhibition of parasite progression, highly synchronized ring stage parasites were treated with 10 µM DMS and their development were monitored at different time intervals (0, 18, 34, 45 hpi) by preparing thin blood smears of the treated and untreated infected erythrocytes. As observed in the blood smears, DMS drastically affected progression of the parasite from rings to trophozoites, stalling the parasite progress at the ring stage as compared to the untreated parasite infected control. Halt in the parasite growth could be accounted to the formation of ‘pyknotic body’ after 36 hours of the treatment (Fig. 2b). This progression arrest coincided with a significant drop in the S1P levels of the erythrocytes after DMS treatment. These findings strongly support the hypothesis, which suggested that SphK-1 mediated production of S1P plays an important role in *Plasmodium* growth and development in the erythrocytes. Further, to ascertain whether the host SphK-1 inhibition mediated retardation of parasite growth and progression in independent of DMS-induced topological aberration in host membrane, we have performed SEM-based imaging of both infected and uninfected erythrocytes in presence of DMS. The SEM micrographs hardly show any significant changes in infected erythrocytes, following DMS treatment as compared to the respective untreated control (Fig. 2c). Similar observation could also be inferred from the experiments involving uninfected erythrocytes treated with DMS (Fig. 2c).

**Figure 2.**
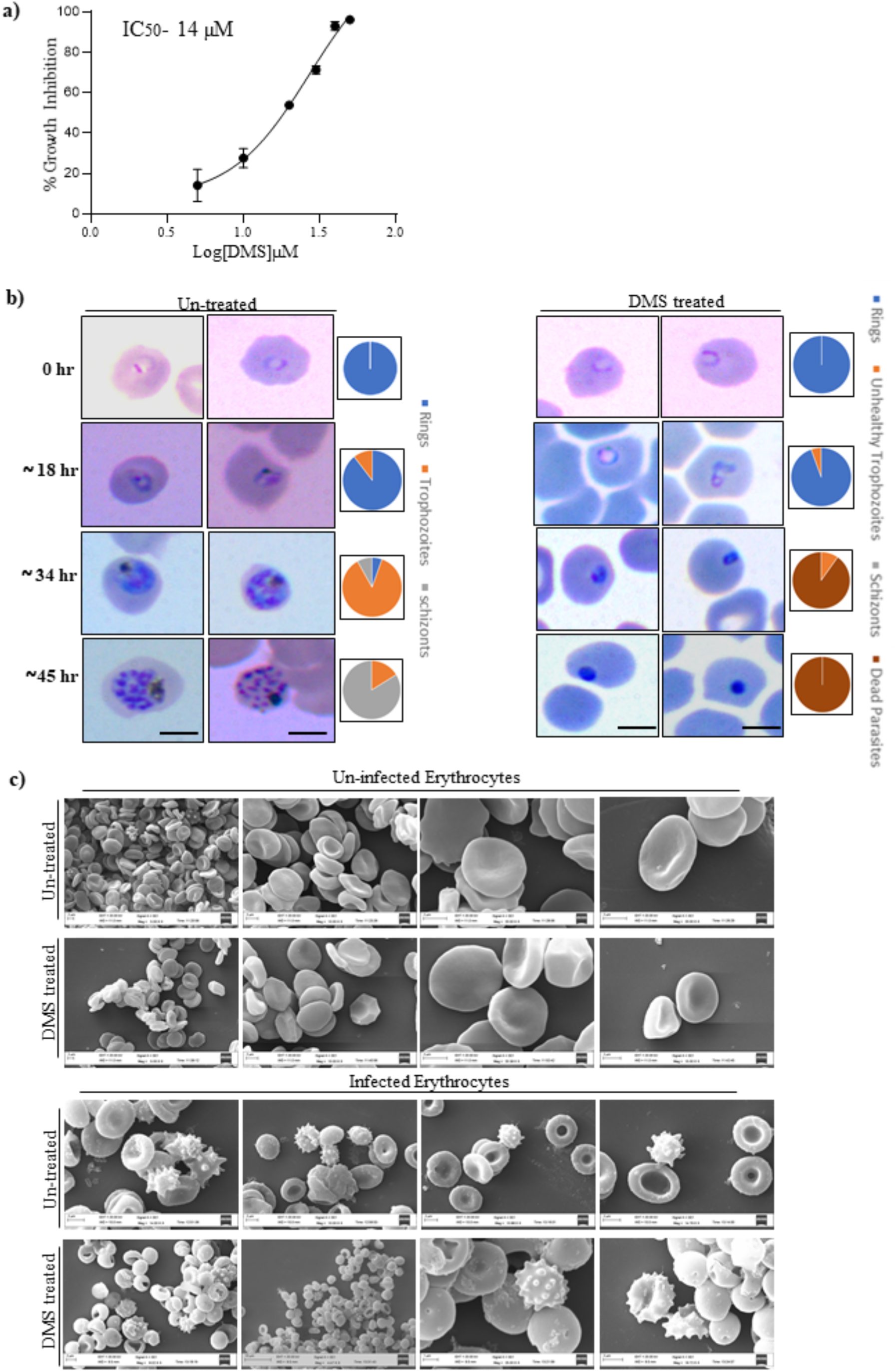
Inhibition of host SphK-1 blocks parasite growth and progression. **A**) Percentage inhibition of *P. falciparum* growth was evaluated at different DMS concentrations (0-40 µM), as presented in the bar graph. Three independent experiments were performed in triplicates with 96-well plates using a SYBR-green assay. The IC_50_ value was determined as 14 µM for the parasite growth inhibition with DMS. **B**) Visualization of stage specific inhibition of *P. falciparum* progression following DMS treatment was depicted by light microscopic images of Giemsa stained ring, trophozoite and schizont stages. **C**) Scanning electron micrographs of DMS-treated infected and uninfected erythrocytes represented comparative morphometric analysis. Scale bars = 5 µ m

### Inhibition of erythrocyte SphK-1 results in decrease in lactate production

The intra-erythrocytic stages of *P. falciparum* lack a functional citric acid cycle and is largely reliant on glycolysis to fulfill its very substantial energy requirements[36],[37]. Erythrocytes infected with mature trophozoite stage parasites consume glucose up to two folds of magnitude faster than uninfected erythrocytes, eventually converting it to lactic acid[10],[11],[38]. To dissect this mechanism of SphK-1 dependent growth progression in parasite infected erythrocytes, and its link to glycolysis, we estimated the amount of lactate formation, an indicator of glycolytic process, in both infected and uninfected erythrocytes in presence of 10 µM DMS for 3 hours. The findings suggested a prominent reduction in lactate production in DMS-treated infected and uninfected erythrocytes, as compared to respective untreated controls (Fig. 3a). Interestingly, parasite stage-specific estimation of lactate levels during intra erythrocytic cycle, represented predominant decrease in lactate production, as clearly evident in rings, trophozoites and schizonts (Fig. 3b), following treatment with 10 µM of DMS for 3 hour. These data depict that; lowering of S1P levels by DMS-based inhibition of host SphK1 can lead to compromised glycolysis, which is manifested as reduced lactate levels in intra-erythrocytic cycle of *P. falciparum*.

**Figure 3.**
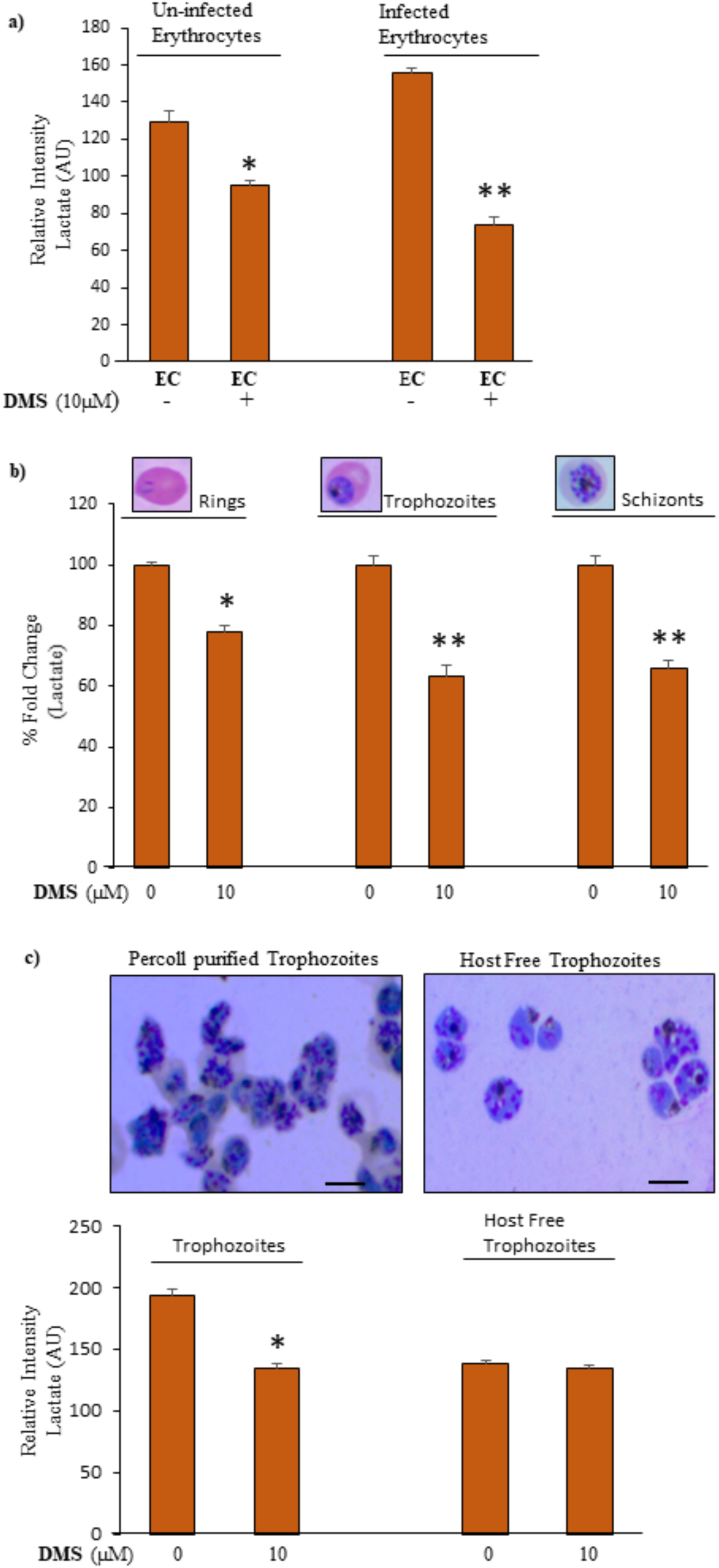
Inhibition of host SphK-1 results in reduction of lactate production. **A)** Quantification of lactate in EC microenvironments of uninfected and infected erythrocytes treated with DMS (10 µM), was represented as changes in relative intensities. **B)** Evaluation of lactate levels in asexual blood stages following DMS treatment, was plotted as percentage fold change. **C)** Light microscopic images of saponized/host-free trophozoites and percoll purified trophozoites were visually compared following Giemsa-staining. Following identification, the purified and saponized samples were further used for above experiment. Representative bar graph displayed changes in lactate levels as relative intensities for trophozoites and saponized/host-free parasites

To gain further insight, whether the observed metabolic alterations after inhibitor treatment were mediated specifically through erythrocytes or it directly impacted the parasite, we have elucidated the changes in lactate levels in host-free parasite isolated by saponin lysis, in response to the inhibitor. To determine the intrinsic lactate level of the trophozoite stage along with the saponin lysed host-free parasites were treated with the inhibitor for 2 hours. Interestingly, the lactate levels were found to be lowered as compared to their respective untreated controls except in the case of saponin lysed parasites wherein, there was no significant difference in the lactate levels of DMS treated and untreated parasites could be detected (Fig. 3c). Lowering of lactate levels could be accountable to either down regulated glycolysis rate and/or reduced glucose uptake in trophozoite infected and uninfected erythrocytes. However, saponin lysed/host-free DMS treated trophozoites demonstrated negligible difference in the lactate levels as compared to the untreated control, suggesting erythrocytes as the mediator of S1P-dependent glycolysis not the parasite (Fig. 3c). Altogether, these readouts suggested that host SphK-1 contributes to the regulation of glycolysis and its effect on parasite growth is mediated solely through the erythrocyte.

### Abrogated S1P synthesis results in reduced glucose uptake

Further, we investigated the role of S1P in glucose uptake and glycolysis, as erythrocytes are solely dependent on glycolysis to meet their energy requirements. Towards this, we measured the glucose uptake in parasite-infected erythrocytes via fluorescence labelling and fluorescence-activated cell sorting (FACS) using a non-metabolizable glucose analog, 2-NBDG, which is fluorescently tagged. 2-NBDG accumulates inside the cells by entering through the glucose transporters but it does not enter into the glycolysis pathway[38],[39],[40]. Parasite infected erythrocytes were pre-incubated with DMS or solvent control for 3 hours at 37°C. The cells were then incubated with fluorescently labeled 2-NBDG and its uptake was measured after washing the cells with PBS using confocal microscopy and flow cytometry. Live-cell imaging of DMS-treated infected erythrocytes represented drastic reduction in glucose uptake, as represented by diminished green fluorescence indicating lower uptake of 2-NBDG. While, the untreated infected erythrocytes displayed prominent green fluorescence suggesting healthy uptake of glucose analogues (Fig. 4a). The fluorescence intensities were plotted using Image J for the treated and untreated cells indicating the same (Fig. 4a). To confirm whether the impaired uptake of glucose analogue in DMS-treated infected erythrocytes was due to dysregulated glycolysis but not compromised cell viability, we used a strategy using dual dye-based assay. Regarding this, we have used Syto9 in combination with Propidium Iodide (Thermo Fisher Scientific, Waltham, Massachusetts, US) to delineate viability in DMS-treated infected erythrocytes. The finding revealed that the viability was uncompromised in DMS-treated infected erythrocytes with significant reduction in 2-NBDG uptake, suggesting the impairment in glucose uptake was mainly due to altered S1P level following host SphK-1 inhibition by DMS. Further, flow cytometry analysis of 2-NBDG uptake in DMS-treated infected erythrocytes demonstrated a 10-fold shift in fluorescence intensity indicating enhanced uptake of 2-NBDG, whereas, the untreated infected erythrocytes showed severe depletion in 2-NBDG uptake, as shown by diminished intensity peak in representative histogram (Fig. 4b). These results suggested that the uptake of glucose analogue, a signature of healthy glycolysis, was strongly repressed by the host SphK-1 inhibition as compared to their respective controls.

**Figure 4.**
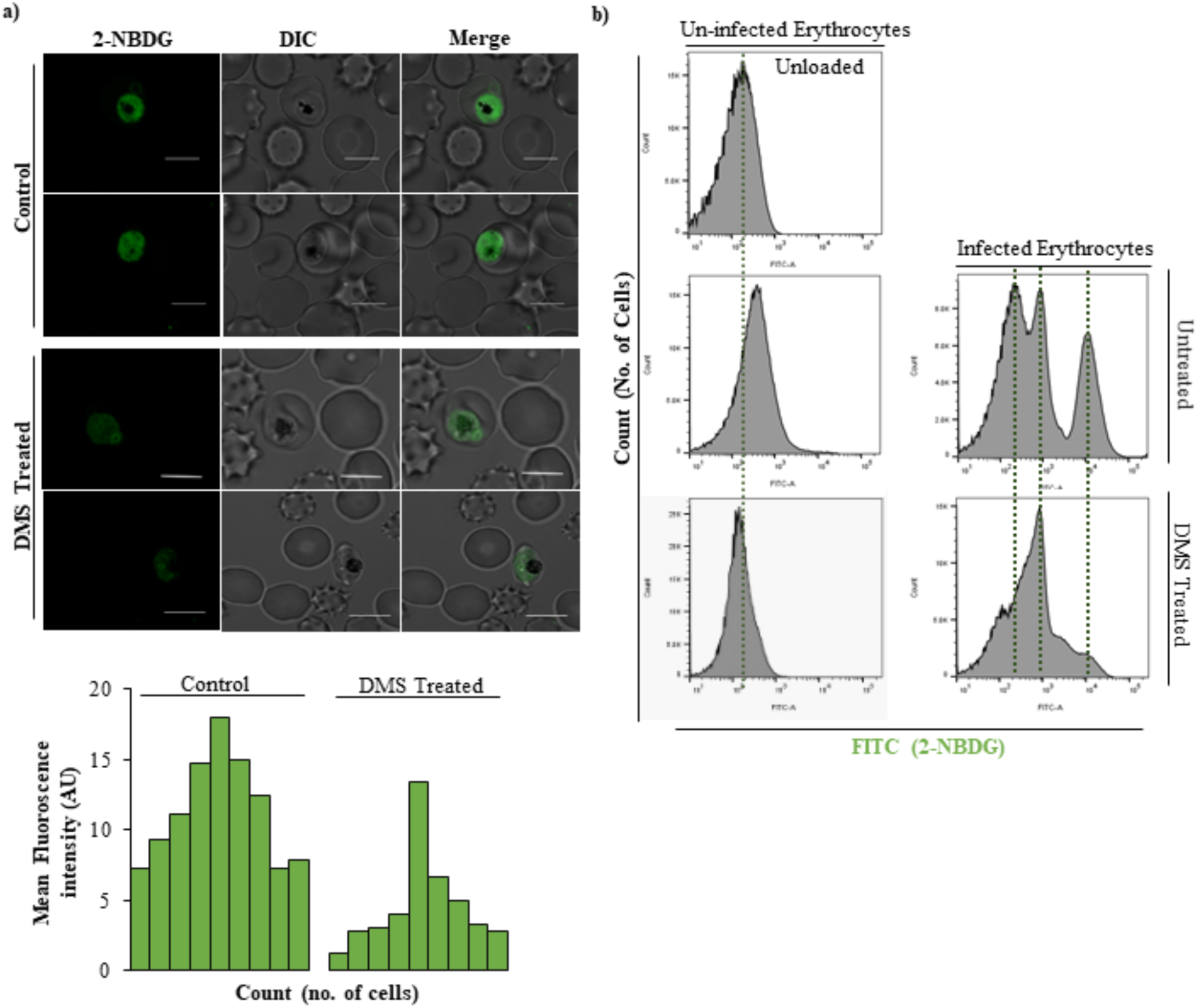
Detection of glucose uptake by parasite infected erythrocytes upon host SphK-1 inhibition. **A)** Evaluation of 2-NBDG uptake in parasite infected erythrocytes following DMS treatment by live-cell imaging. Respective MFI of individual infected erythrocytes were plotted against individual untreated cells. **B**) Representative histograms depict changes in number of FITC^positive^ population in flow cytometry analysis correlating to 2-NBDG uptake following DMS treatment in infected erythrocytes.

### Inhibition of erythrocyte SphK-1 leads to change in translocation of glycolytic enzyme GAPDH from membrane-to-cytosol

Intracellular S1P in erythrocytes has been known to release membrane-bound GAPDH, a glycolytic enzyme to cytosol in response to hypoxia[17]. Since, host SphK-1 inhibition led to depletion in S1P levels in infected erythrocytes, we assumed translocation of GAPDH from membrane to cytosol might be also be hampered in DMS-treated infected erythrocytes. To validate the same, we checked the levels of GAPDH in cytosolic and membrane fractions of parasite infected and uninfected erythrocytes. The GAPDH level was detected by immunoblotting and immunolabelling. The total cell lysate was used as a control. After DMS treatment, both the membrane and cytosolic fractions were separated for probing with anti-GAPDH mouse antibody. As expected, DMS treatment restricted the translocation of GAPDH to the erythrocyte membrane, as evident in immunoblot and its respective band intensities in both membrane and cytosolic fractions (Fig. 5a). These findings confirmed low release of GAPDH to the cytosol due to altered glycolysis in DMS-treated infected erythrocytes. Further, we visualized the localization of GAPDH in the cells by immunolabelling using confocal imaging. After DMS treatment, GAPDH was mainly restricted to the erythrocytic membrane rather than translocating to the cytosol, suggesting decreased glycolysis, which can be corroborated to our previous findings (Fig. 5b). Collectively, these results suggested low S1P levels in DMS-treated infected erythrocytes can lead to aborted glycolysis.

**Figure 5.**
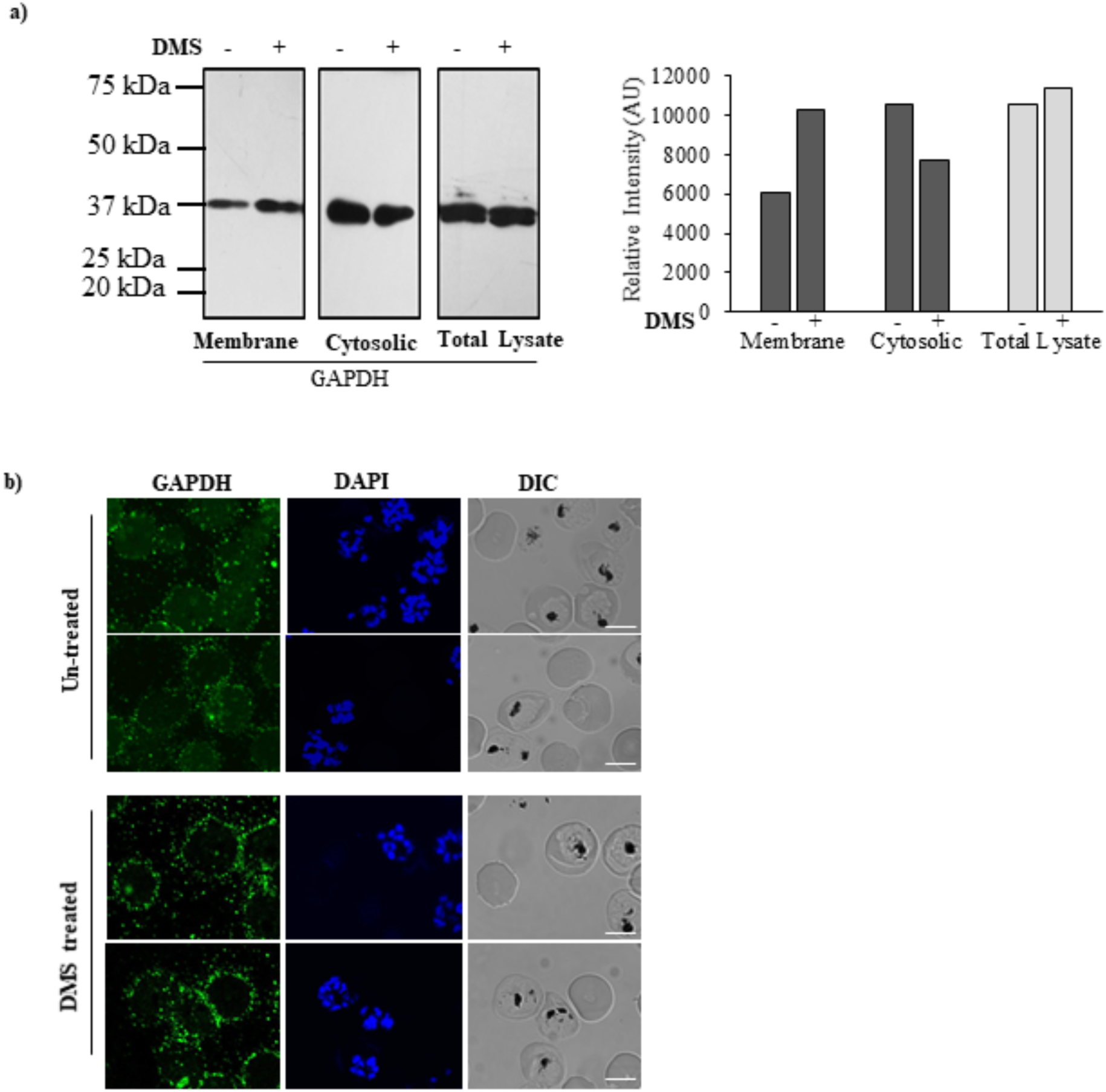
Host SphK-1 mediated translocation of GAPDH in parasite infected erythrocytes. **A)** Detection of localized level of GAPDH in both membrane and cytosolic fractions of DMS-treated infected erythrocytes. Total cell lysate was used as a loading control. Change in the level was plotted as relative intensities of bands detected in different cellular fractions. **B**) Confocal micrographs demonstrate GAPDH level and localization in both DMS-treated and untreated infected erythrocytes.

## Discussion

It is noteworthy that, circulating erythrocytes demonstrate high storage of S1P, a crucial signaling biolipid, as compared to other peripheral tissues, especially due to lack of S1P degrading enzyme, Sphingosine 1 phosphate lyase (SPL)[17]. However, the erythrocytes do express S1P transporter Mfsd2b[31] and the concentration gradient of S1P from circulating erythrocytes to peripheral tissues governs several critical physiological processes, including vascular integrity, trafficking of lymphocytes and bone homeostasis *etc*[41],[20],[42].

S1P also play as a rheostat for maintaining balance between cytostasis and apoptosis[43]. In addition, the SphK-1/S1P signaling nexus contributes to the development and progression of various diseases including, Huntington’s disease [44] and ulcerative colitis[45] *etc*. Thus, modulation of SphK-1 enzyme and its products have become the prime target in order to reduce the disease severity[46],[47]. S1P is catalyzed by two isoforms of Sphingosine kinase, SphK (isoforms SphK-1 and SphK-2), and can turn on various cellular processes by activating a family of G protein-coupled receptors, sphingosine-1-phosphate receptor 1-5 (S1P_1-5_)[48].

Accumulating body of evidences reveal strong role of deregulated S1P metabolism in parasite-born diseases, including trypanosomiasis, leishmaniasis and cerebral malaria[49],[50],[51]. However, studies involving host-SphK1 mediated manipulation of parasite growth and progression are still at their infancy and unraveling the same would provide a breakthrough to present alternative targets for the drug therapies. With this perspective, we have delineated the possible role of host-SphK1 activity on growth and progression of *P. falciparum*, the causative agent of malaria, one of the deadly parasitic diseases.

To achieve this, we targeted host SphK-1 activity by using a specific inhibitor DMS, which completely depleted the S1P levels in both IC and EC of infected erythrocytes, leading to stalled the parasitic growth and progression with formation of ‘pyknotic bodies’ (Fig. 1, 2). To negate the possibility of DMS-enforced topological alterations in infected erythrocytes as the basis of aborted parasite growth and invasion, we have performed SEM-based analysis of both uninfected and infected erythrocyte membranes after DMS treatment. To emphasize, there was no significant changes could be identified in membrane structures of infected erythrocytes even after DMS treatment (Fig. 2c). Similar inference was drawn from the experiment involving uninfected erythrocytes with DMS (Fig. 2c). Since erythrocytes mainly harbor SphK-1, but no S1P receptor for its innate signaling, we hypothesized that inhibiting host-SphK1, might be attenuating the host-S1P dependent parasite survival mechanism as well. During *Plasmodium* infection, erythrocytes demonstrate 6-fold increase in phospholipids (PL), along with a sharp rise in glycolytic flux, with glucose uptake upto 50 folds, predominant in metabolically progressive stages such as (trophozoite and schizont)[52],[53],[54] suggesting host-dependency of the parasite for fulfilling its metabolic needs[55].

To understand, whether inhibition of host-SphK1 during infection might block glycolysis in metabolically active stages of *P. falciparum*, mainly in trophozoite and schizont; we have estimated the levels of lactate, a signature metabolite in the same following DMS treatment. The results depict, drastic switch of metabolically active stage of parasites to metabolically dormant state, leading to parasite growth retardation (Fig. 2, 3). This data strongly advocated the role of host-SphK1 in regulation of glycolysis-dependent growth and progression of parasites. Further, to confirm the hypothesis, which suggested that altered glycolysis of intra-erythrocytic cycles is mediated specifically through erythrocytes not via the parasites; we also estimated the lactate levels in DMS-treated saponized/host-free parasites, as a proof-of-concept. The observation clearly ruled out the involvement of parasite-mediated glycolysis, as no change in lactate levels could be detected in host-free parasites (Fig. 3). To elucidate the impact of host SphK-1 inhibition on glucose uptake, during intra-erythrocytic development, we have measured the live uptake of glucose using a fluorescent-labeled glucose analogue (2-NBDG). The results presented abrogated glucose uptake in DMS-treated infected erythrocytes, imposing a direct role of host-SphK-1 in regulation of glycolysis (Fig. 4).

According to a recent study by K Sun et al., SphK-1 activity gets elevated in erythrocytes under hypoxic conditions, leading to enhanced S1P level and thereby, increasing its binding to deoxygenated hemoglobin (deoxy-Hb). Subsequently, this facilitates deoxy-Hb anchorage to the membrane, leading to more release of membrane-bound GAPDH to the cytosol which then increases the erythrocytic glycolysis[17]. To further evaluate, whether abolishing the host-SphK-1 activity in DMS-treated infected erythrocytes, would dysregulate the GAPDH level, we have evaluated the level of GAPDH both in membrane and cytosolic fractions. The findings unraveled reduced translocation of GAPDH from membrane-to-cytosol in DMS-treated infected erythrocytes, as evident from altered localized level of GAPDH, in both immunoblot and confocal micrographs (Fig. 5). To summarize, our study highlights two important findings; firstly, the inhibition of host SphK-1 activity can abrogate intra-erythrocytic growth and progression of *P. falciparum*, and secondly, diminished S1P levels can lead to host-mediated altered glycolysis resulting in growth retardation of parasites (Fig. 6). Overall, this study introduces SphK-1/S1P signaling nexus in erythrocytes as the alternate pathway for *P. falciparum* survival. Since, knocking down of host SphK-1 is not lethal, thus targeting the same would eliminate the problem of resistance. Conceivably, further elucidation of SphK-1/S1P signaling pathways during parasite infection, might aid in developing novel anti-malarial chemotherapeutics.

**Figure 6.**
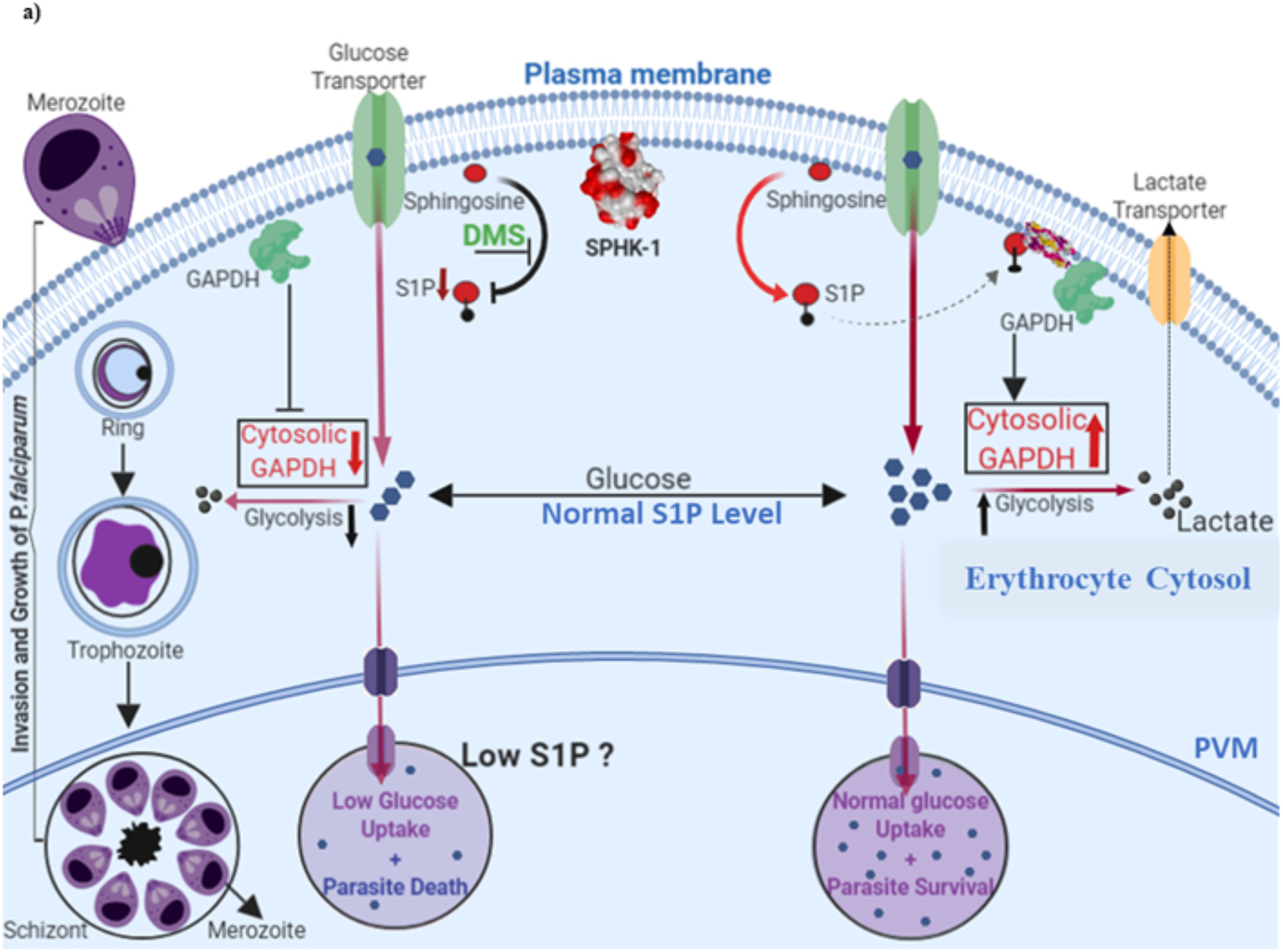
In the proposed working model, life cycle of blood stage parasite is shown. In normal condition, S1P bind to deoxy-Hb and facilitates binding of deoxy-Hb to membrane and release of GAPDH; increased cytosolic GAPDH accelerates glycolysis and generate lactate which is a by-product of glycolysis and does not stall the growth of parasite. In case of DMS-mediated inhibition of host SphK-1, S1P level gets reduced, leading to altered binding with deoxy-Hb. Thus, it does not facilitate GAPDH to cytosol due to which glycolysis is suppressed leading to retarded parasite growth.

## Acknowledgments

We are thankful to Advanced Instrumentation and Research Facility (AIRF), Jawaharlal Nehru University (JNU), New Delhi for Central Instrumentation Facility (CIF) of Special Centre for Molecular Medicine, JNU for confocal microscopy, LC/MS analysis and other instruments facilities. This work was supported by Bio-Scientist Award and Innovative Young Biotechnologist Award (IYBA) from the Department of Biotechnology, Ministry of Science and Technology, Government of India (DBT). Shailja Singh is a recipient of the National Bio scientist and IYBA Award from DBT. R.K.S. is supported by CSIR-UGC fellowship. M.S. is financially supported by Shiv Nadar Foundation fellowships. We acknowledge the financial support from Science and Engineering Research Board (SERB, File no. EMR/2016/005644), India, National Institutes of Health (NIH, Grant no. U19AI089676-09), United States, Department of Science & Technology-Promotion of University Research and Scientific Excellence (DST-PURSE, Phase II, JNU), India. The funders had no role in study design, data collection and analysis, decision to publish, or preparation of the manuscript.

## Author Contributions

SS conceived and designed the research. RKS performed research and SS, RKS & SP analyzed the data. RKS and SS conducted the lipid extraction and estimation experiments. MS and RKS performed microscopy experiments. SS, RKS, MS & SP wrote the manuscript. SS finally edited the manuscript.

