## Supplementary Material for "*Plasmodium falciparum* growth is regulated by Sphingosine 1 phosphate produced by Host Erythrocyte Membrane Sphingosine kinase 1"

a

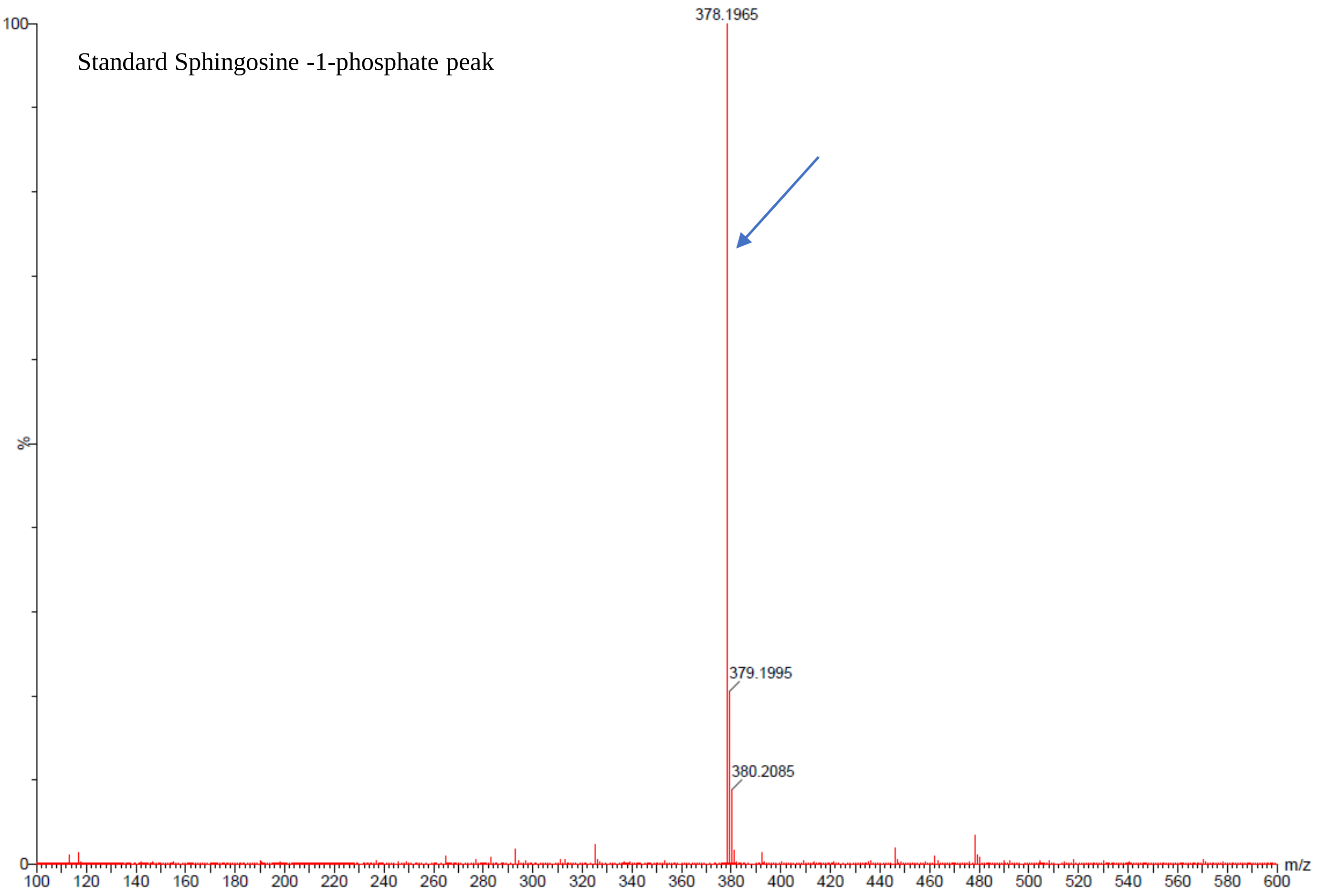


b

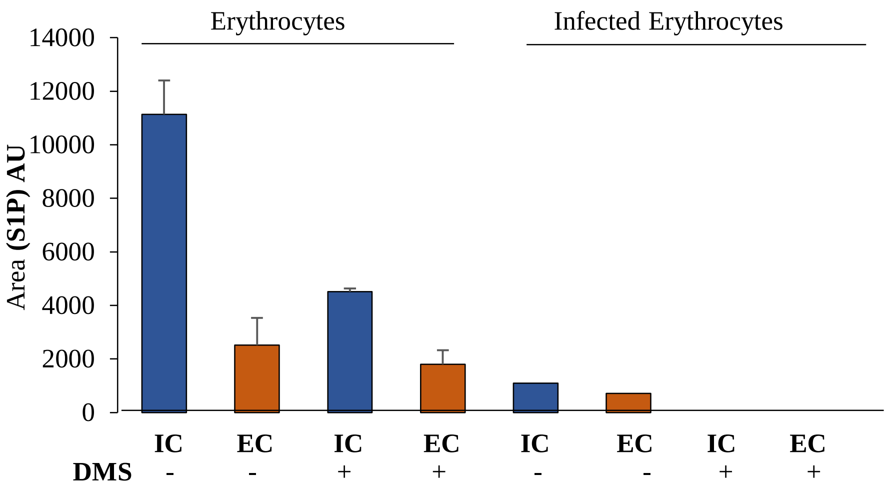


**Supplementary Figure 1. Characteristic peak of S1P in MS spectra**. S1P in methanol was subjected to LC/MS analysis and generated a characteristic peak at position 378.16 acquired in MS spectra **(A)**. Representative bar graphs display LC/MS-based quantification of intracellular (IC) and extracellular (EC) S1P levels in parasite infected and uninfected erythrocytes in presence of DMS (10 µM). Acquired S1P-specific peak highlighted in the MS spectra **(B)**.

a)

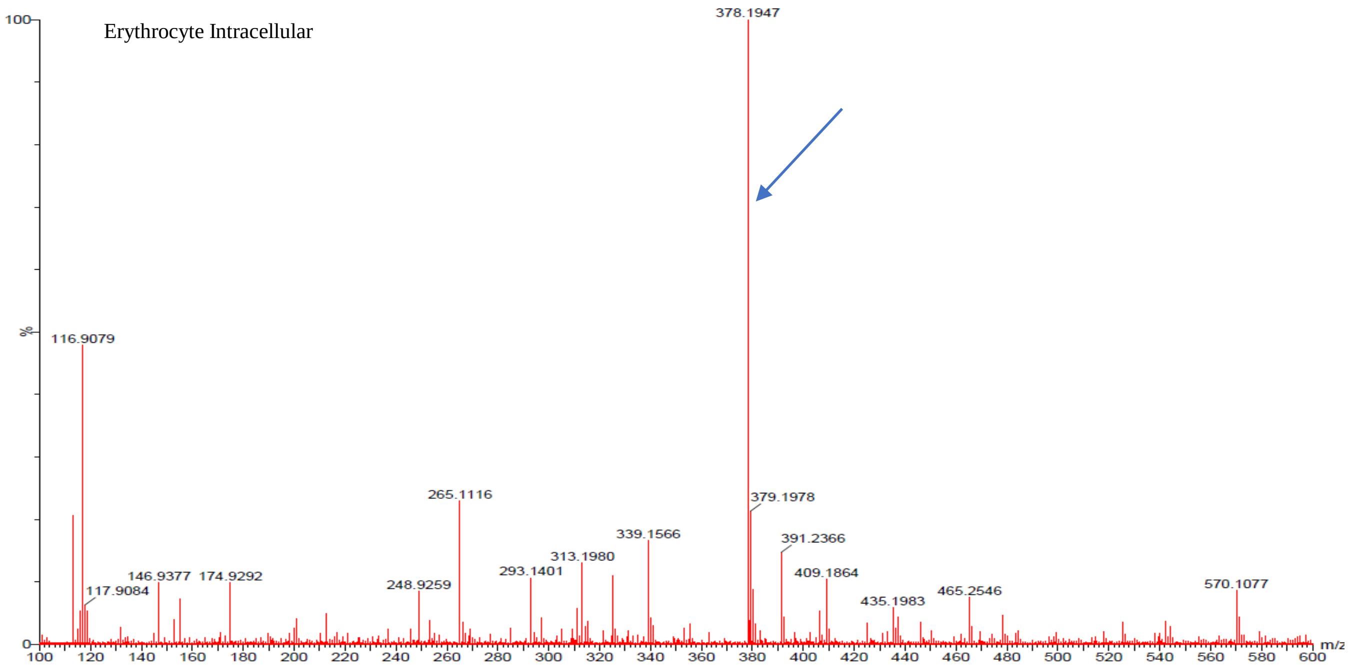
Erythrocyte Intracellular
378.1947
116.9079
265.1116
379.1978
339.1566
391.2366
313.1980
293.1401
409.1864
146.9377
174.9292
248.9259
117.9084
435.1983
465.2546
570.1077
%
m/z

b)

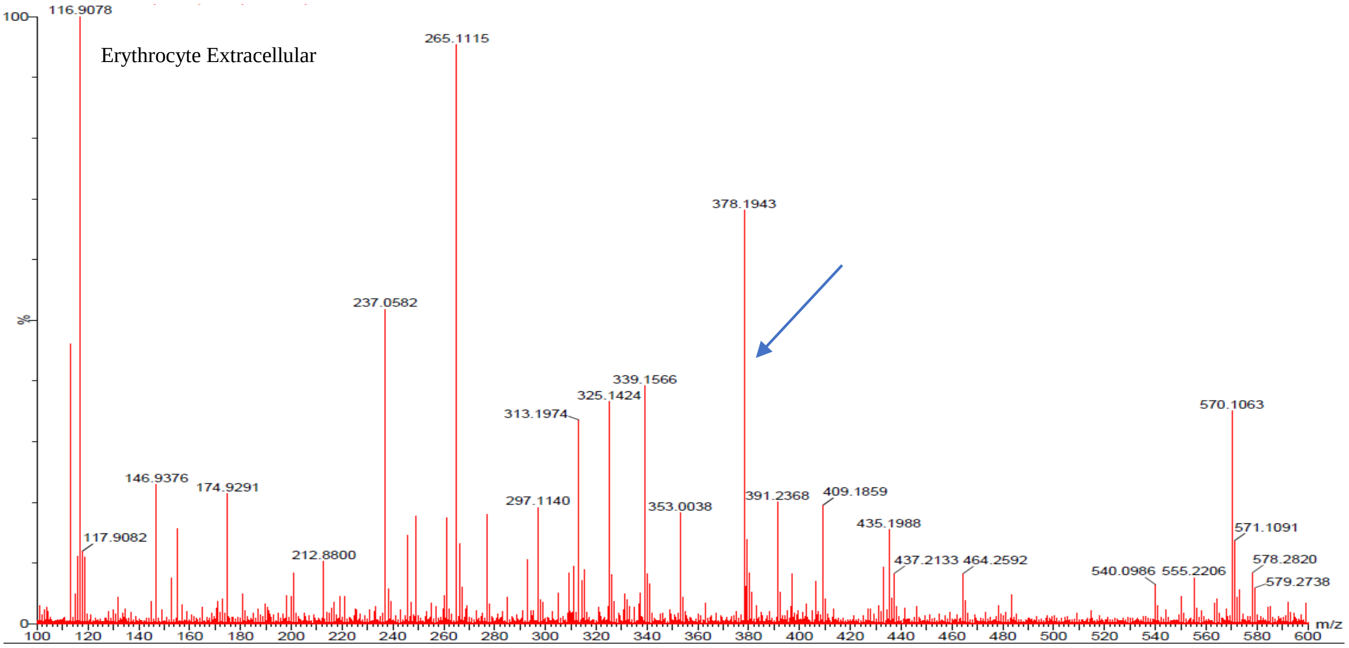
Erythrocyte Extracellular
116.9078
265.1115
378.1943
237.0582
339.1566
325.1424
313.1974
570.1063
146.9376
174.9291
297.1140
391.2368
409.1859
353.0038
435.1988
117.9082
212.8800
571.1091
437.2133
464.2592
578.2820
540.0986
555.2206
579.2738
%
m/z

c)

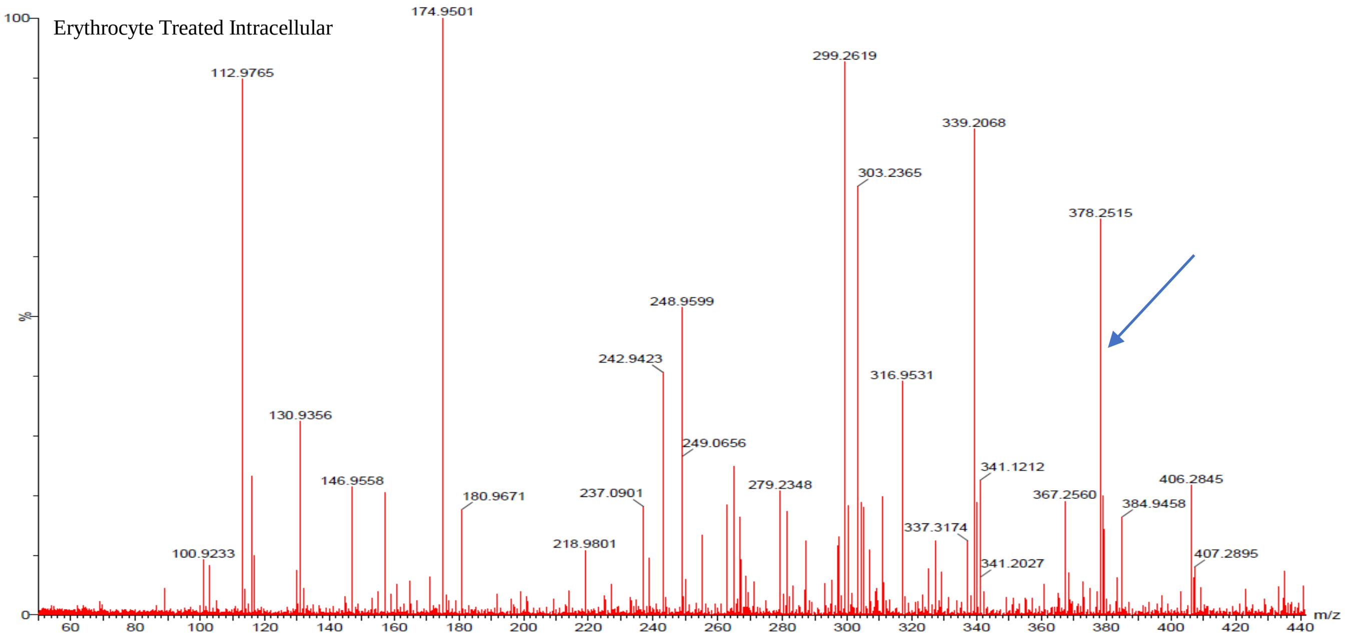
Erythrocyte Treated Intracellular
174.9501
112.9765
299.2619
339.2068
303.2365
378.2515
248.9599
242.9423
316.9531
130.9356
249.0656
341.1212
406.2845
146.9558
180.9671
237.0901
279.2348
367.2560
384.9458
337.3174
218.9801
100.9233
341.2027
407.2895
%
m/z

d)

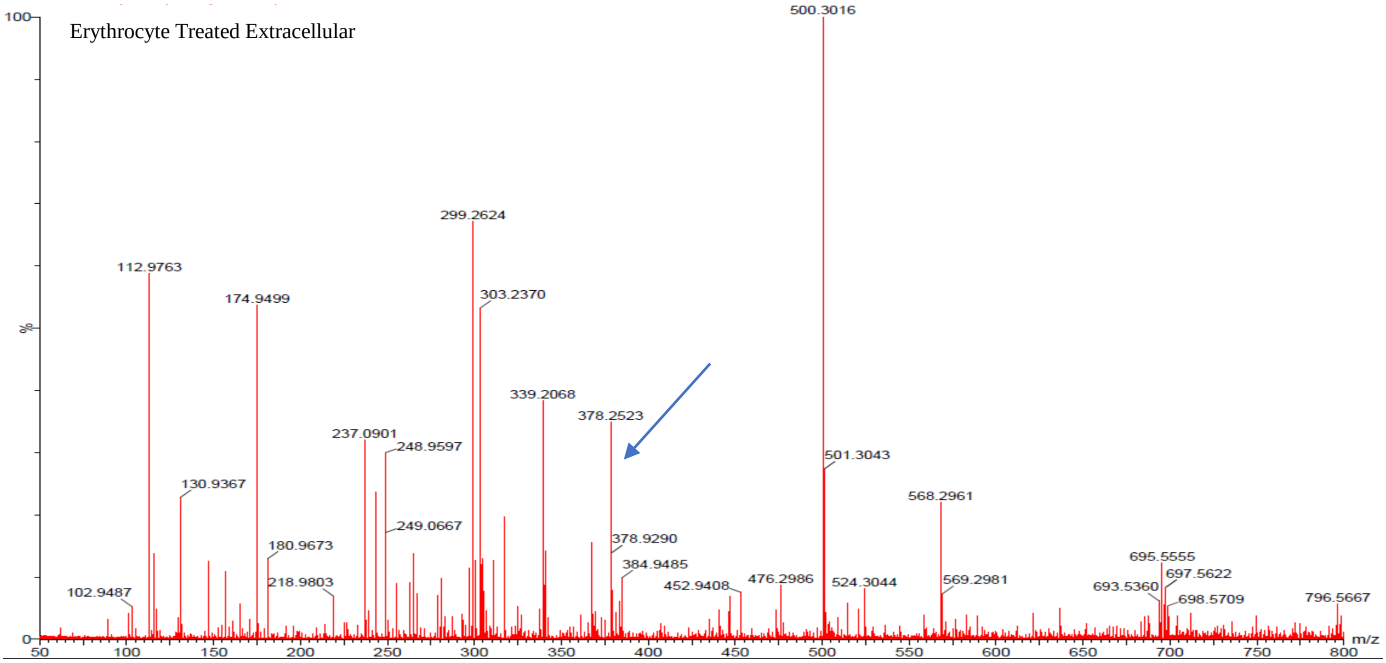
Erythrocyte Treated Extracellular
500.3016
299.2624
112.9763
174.9499
303.2370
339.2068
378.2523
237.0901
248.9597
501.3043
130.9367
568.2961
249.0667
180.9673
378.9290
384.9485
695.5555
697.5622
218.9803
476.2986
452.9408
524.3044
569.2981
693.5360
102.9487
698.5709
796.5667
%
m/z

e)

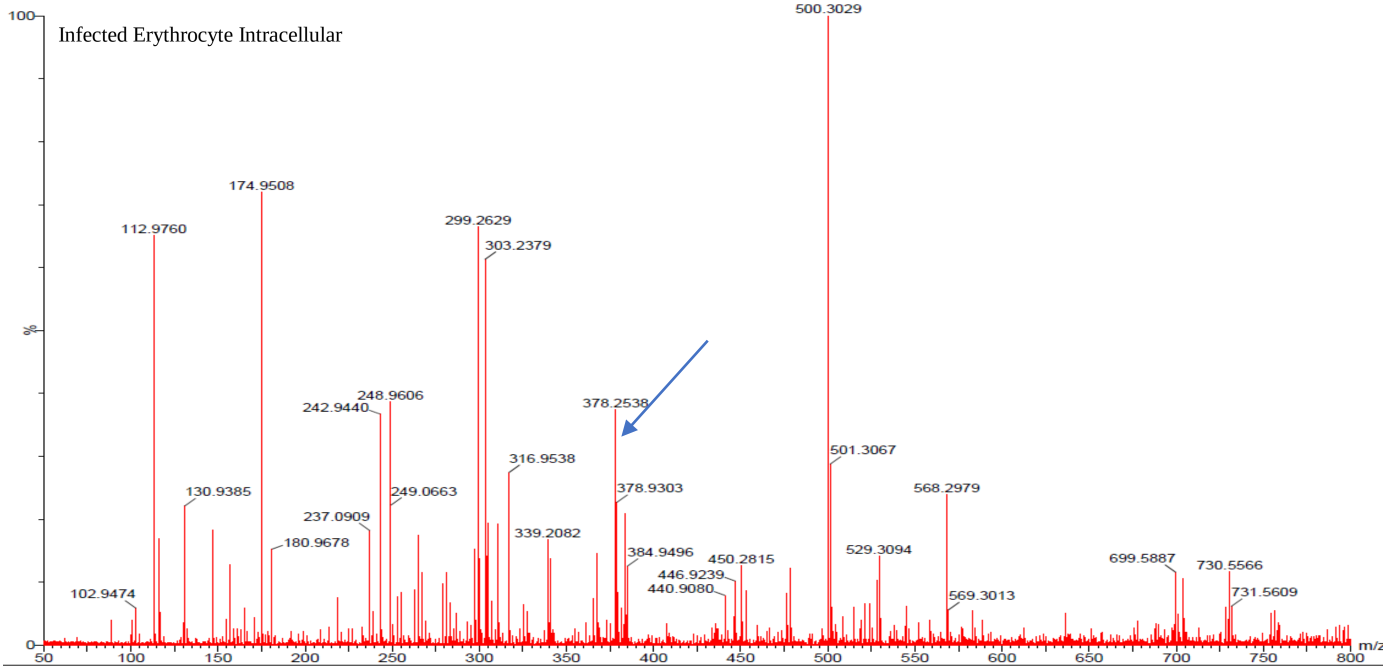
Infected Erythrocyte Intracellular
500.3029
174.9508
112.9760
299.2629
303.2379
248.9606
242.9440
378.2538
501.3067
316.9538
130.9385
249.0663
378.9303
568.2979
237.0909
180.9678
339.2082
384.9496
529.3094
450.2815
699.5887
730.5566
446.9239
440.9080
102.9474
569.3013
731.5609
%
m/z

f)

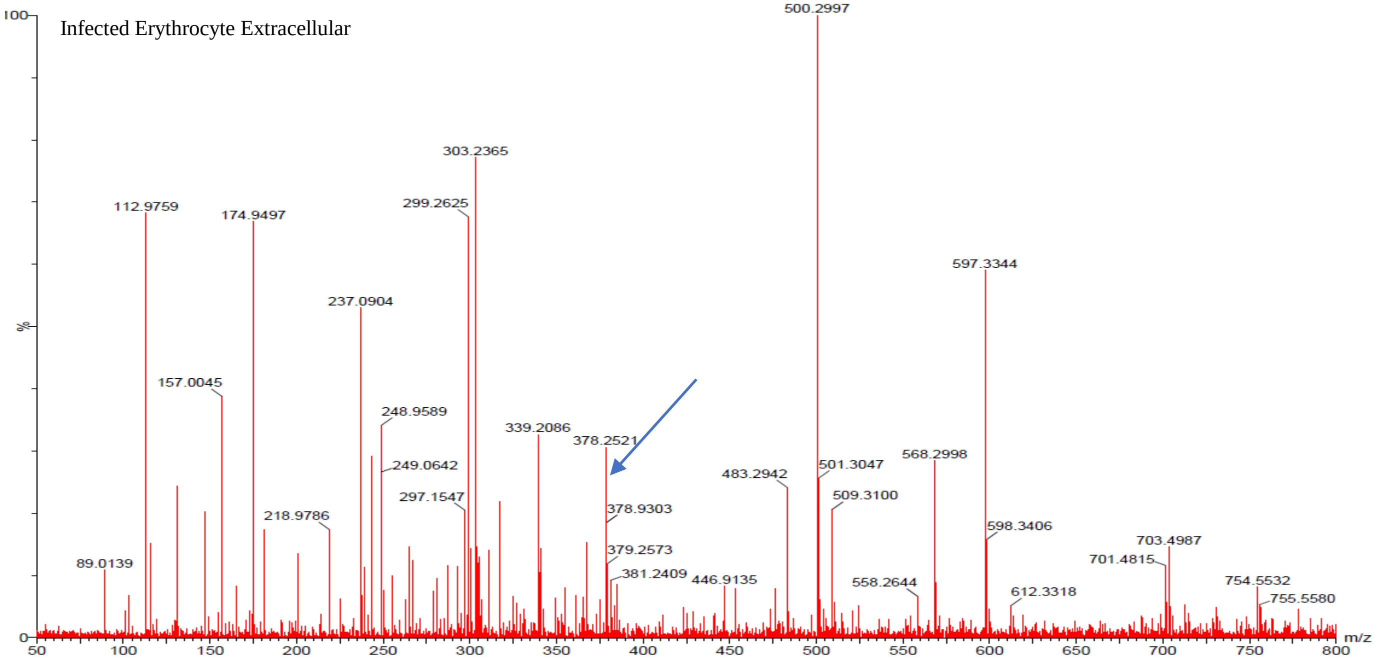
Infected Erythrocyte Extracellular
500.2997
303.2365
112.9759
299.2625
174.9497
597.3344
237.0904
157.0045
248.9589
339.2086
378.2521
568.2998
249.0642
483.2942
501.3047
297.1547
509.3100
378.9303
218.9786
598.3406
703.4987
379.2573
89.0139
701.4815
381.2409
446.9135
558.2644
754.5532
612.3318
755.5580
%
m/z

g)

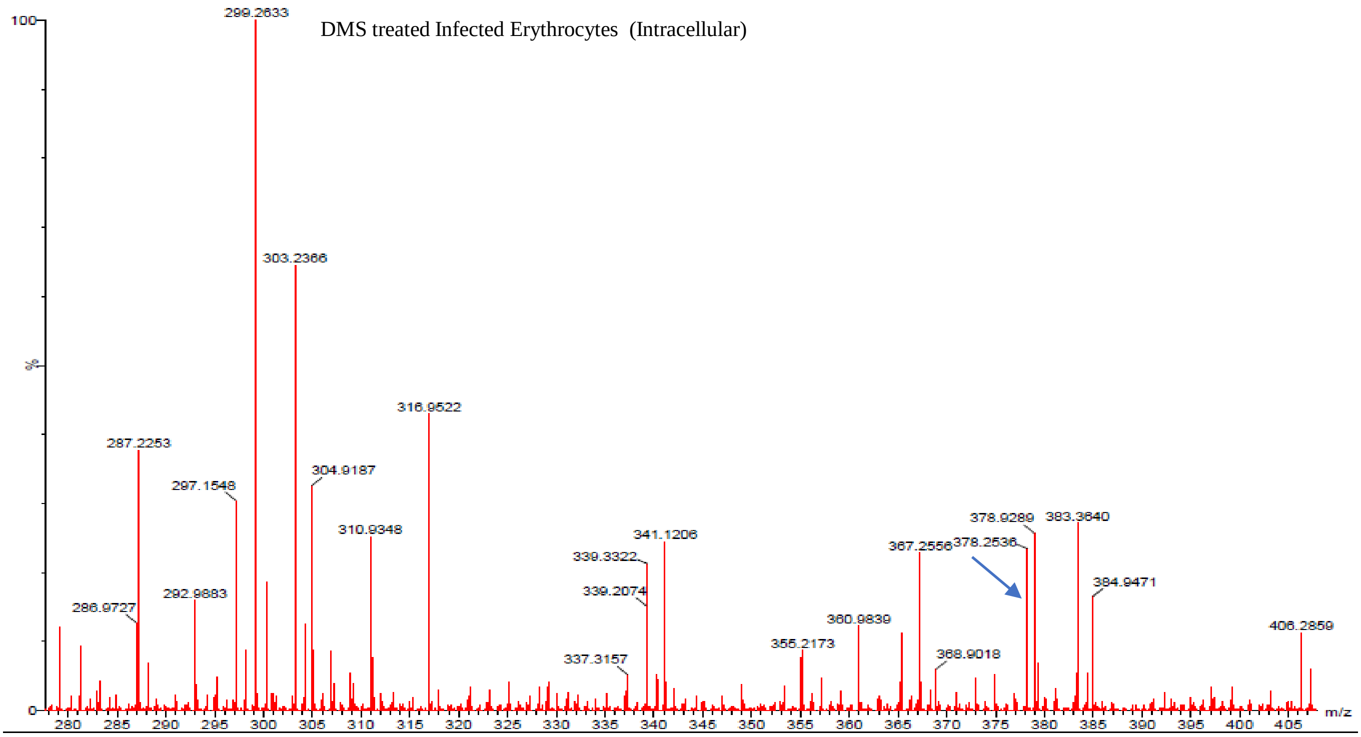


h)

Infected Erythrocyte Treated Extracellular Not Detected(Because of very low level)

**Supplementary Figure 2. S1P detection in IC and EC milieu of uninfected erythrocytes and infected erythrocyte with DMS treatment and without treatment.** Lipids were extracted from supernatant and lysed cells from uninfected erythrocytes and infected erythrocyte after DMS treatment and without treatment. The extracted lipids were subjected to LC/MS analysis for S1P detection. S1P characteristic peaks were detected at position 378.25 in the MS spectra for IC as well as EC milieu **(A-G)**. In **Figure 2 (H)** Due to lower level of S1P in DMS treated cells, peak was not detected.
